# Adaptive radioresistance of Enterohemorrhagic *Escherichia coli* O157:H7 results in genomic loss of shiga toxin-encoding prophages

**DOI:** 10.1101/2022.08.08.503262

**Authors:** Ghizlane Gaougaou, Antony T. Vincent, Kateryna Krylova, Hajer Habouria, Hicham Bessaiah, Amina Baraketi, Frédéric J. Veyrier, Charles M. Dozois, Eric Déziel, Monique Lacroix

## Abstract

Enterohemorragic *Escherichia coli* (EHEC) O157:H7 is a foodborne pathogen producing shiga toxins (Stx1 and Stx2), can cause hemorrhagic diarrhea, and life-threatening infections. O157:H7 strain EDL933 carries prophages CP-933V and BP-933W that encode shiga toxin genes (*stx1* and *stx2* respectively). The aim of this work was to investigate the mechanisms of adaptive resistance of EHEC strain EDL933 to a typically “lethal” dose of γ-irradiation (1.5 kGy). Adaptive selection through six passages of exposure to 1.5 kGy resulted in the loss of CP-933V and BP-933W prophages from the genome and mutations within three genes: *wrbA, rpoA*, and Wt_02639 (*molY*). Three selected EHEC clones that became irradiation-adapted to the 1.5 kGy dose (C1, C2 and C3) demonstrated increased resistance to oxidative stress, sensitivity to acid pH, and decreased cytotoxicity to Vero cells. To confirm that loss of prophages plays a role in increased radioresistance, C1 and C2 clones were exposed to bacteriophage containing lysates. Although, phage BP-933W could lysogenize C1, C2, and *E*.*coli* K-12 strain MG1655, it was not found to have integrated into the bacterial chromosome in C1-Φ and C2-Φ lysogens. Interestingly, for the *E. coli* K-12 lysogen (K12-Φ), BP-933W DNA had integrated at the *wrbA* gene (K12-Φ). Both C1-Φ and C2-Φ lysogens regained sensitivity to oxidative stress, were more effectively killed by a 1.5 kGy γ-irradiation dose and had regained cytotoxicity and acid resistance phenotypes. Further, the K12-Φ lysogen became cytotoxic, more sensitive to γ-irradiation and oxidative stress and slightly more acid resistant.

**Importance:** Gamma (γ)-irradiation of food products can provide an effective means of eliminating bacterial pathogens such as enterohemorrhagic *Escherichia coli* (EHEC) O157:H7, a significant foodborne pathogen that can cause severe disease due to the production of Shiga toxins. To decipher the mechanisms of adaptive resistance of the O157:H7 strain EDL933, we evolved clones of this bacterium resistant to a lethal dose of γ-irradiation by repeatedly exposing bacterial cells to irradiation following a growth restoration over six successive passages. Our findings provide evidence that adaptive selection involved modifications in the bacterial genome including deletion of the CP-933V and BP-933W prophages. These mutations in EHAC O157:H7 resulted in loss of *stx1, stx2*, loss of cytotoxicity to epithelial cells and decreased resistance to acidity, critical virulence determinants of EHEC, concomitant with increased resistance to lethal irradiation and oxidative stress. These findings demonstrate that the potential adaptation of EHEC to high doses of radiation would involve elimination of the Stx encoding phages and likely lead to a substantial attenuation of virulence.

## Introduction

Enterohemorragic (EHEC) *Escherichia coli* (*E. coli*) O157:H7 and other Shiga-toxinogenic *E. coli*/Vero-toxinogenic *E. coli* (STEC/VTEC) are an important risk to public health and food safety. EHEC O157:H7 can cause life-threatening diseases that range from mild gastroenteritis to hemorrhagic colitis and, in extreme cases, cause hemolytic uremic syndrome and kidney failure [1, 2]. EHEC O157:H7 is proposed to have evolved from a non-toxigenic *E. coli* O55:H7 strain possessing a *locus of enterocyte effacement* (LEE). This evolution involved four sequential steps: (i) acquisition of *stx2*, (ii) acquisition of plasmid pO157, (iii) acquisition of *stx1*, and (iv) loss of beta-glucuronidase activity and the capacity to ferment D-sorbitol [3].

Bacteriophages are viruses that infect bacteria. They also contribute in several manners to the evolution of their hosts such as by lysogenic conversion and horizontal gene transfer [4], whereby phage genes integrate into bacterial genomes, and can thus increase bacterial gene content. Lysogeny may also result in gain/loss of function and potentially increased fitness or virulence. For example, the virulence of some bacteria rely on bacteriophage-encoded genes acquired through this process [5, 6]. *E. coli* O157:H7 strain EDL933 possesses 1,387 more genes than the non-pathogenic *E. coli* K-12 laboratory strain MG1655. This increased repertoire includes genes encoding alternative metabolic capacities, virulence factors, several phage proteins and hypothetical proteins [7]. *E. coli* 0157:H7 strain EDL933 carries both *stx*1 and *stx*2 genes located on the prophages PV-933V and BP-933W, respectively. These prophages are considered beneficial to their bacterial hosts by providing new functions during lysogenic conversion that can be considered a bacterial-phage co-evolution [8].

Shiga toxin-encoding prophages integrate into the bacterial genome by site-specific recombination through a tyrosine recombinase integrase that binds to the *att* sites to promote recombination and phage integration [9, 10]. The BP-933W prophage is integrated adjacent to the *wrbA* gene involved in the oxidative stress response, and the PV-933V phage is integrated adjacent to the *mlrA* (*yehV)* gene which encodes a regulator involved in curli and cellulose production [11].

To maintain lysogeny, lambdoid prophages constitutively express the *cI* gene that represses expression of genes involved in the lytic cycle. DNA-damaging agents such as UV irradiation and antibiotics, for example mitomycin C, activate the SOS response that leads to a proteolytic cleavage of the CI protein by RecA, resulting in the induction of the lytic cycle [12-16]. The same recombination reaction may promote excision of prophages by excisionase [9, 17]. In some cases, prophages can inactivate the expression of a bacterial gene via their integration into the open reading frame or the adjacent regulatory region [18]. The prophage can also be excised when bacterial cells require this gene function to counter a stress encountered while the bacteriophage is integrated at this site. Active lysogeny is reversible when the excised prophage can remain in the cell as an episome and may reintegrate into the bacterial chromosome in the absence of conditions that favour the lytic cycle [12, 13, 19]. In other cases, excision of the prophage may be followed by phage loss without triggering lytic production or bacterial lysis. DNA damage by exposure to oxidative stress (and mitomycin C) can induce the SOS response and promote a switch from the lysogenic to the lytic states in *E. coli* O157:H7, and can also result in prophage excision [9, 17]. γ-irradiation can also induce the SOS response [14].

γ-irradiation is an established means of prolonging the shelf life of products such as fresh fruits and vegetables, meat, fish and cereals by eliminating pathogens and by reducing the total bacterial count that can lead to food spoilage [20, 21]. γ-irradiation treatments generate an oxidative stress that kills bacterial cells through generation of reactive oxygen species (ROS) from the radiolysis of water. This oxidative stress causes structural damage and physical dysfunction in bacteria: including DNA damage and modifications, disruption of the cellular envelopes; ribosome alterations; and alteration of selective permeability of the membrane [22].

However, many bacteria, including *E. coli* O157:H7, can acquire radioresistance [23]. Herein, we investigated mechanisms underlying how *E. coli* O157:H7 can develop resistance to a normally lethal dose of γ-irradiation and determine what genetic changes can contribute to this process. *E. coli* O157:H7 strain EDL933 carrying both *stx1* and *stx2* was adapted to a 1.5 kGy dose of irradiation, which is normally considered lethal, by several passages. Genome sequencing of the radiation-adapted bacteria and stress resistance studies were performed to determine radiation resistance mechanisms of *E. coli* O157:H7 strain EDL933.

## Results

### Genomic modification of *E. coli* O157:H7 following adaptation to a normally lethal dose of γ-irradiation

To identify adaptations allowing *E. coli* O157:H7 to become more resistant to radiation doses that are normally lethal to *E. coli*, we adapted strain EDL933 to 1.5 kGy irradiation through several passages (**Figure 1A**). Results showed that bacteria were able to survive and grow during 14 days after the first treatment and the adapted bacterial population increased with the number of passages (**Figure 1B**). The viability of 10 clones was tested after the 6^th^ irradiation passage (1.5 kGy). All of them were able to survive directly after this ‘’lethal stress’’ (**Figure 1C**). Since the viability of adapted *E. coli* O157:H7 clones C1, C2 and C3 was the highest (Survival counts of 2.8, 3.71 and 2.4 log CFU/mL following irradiation respectively). The genomes of these three clones were sequenced.

**Figure 1.**
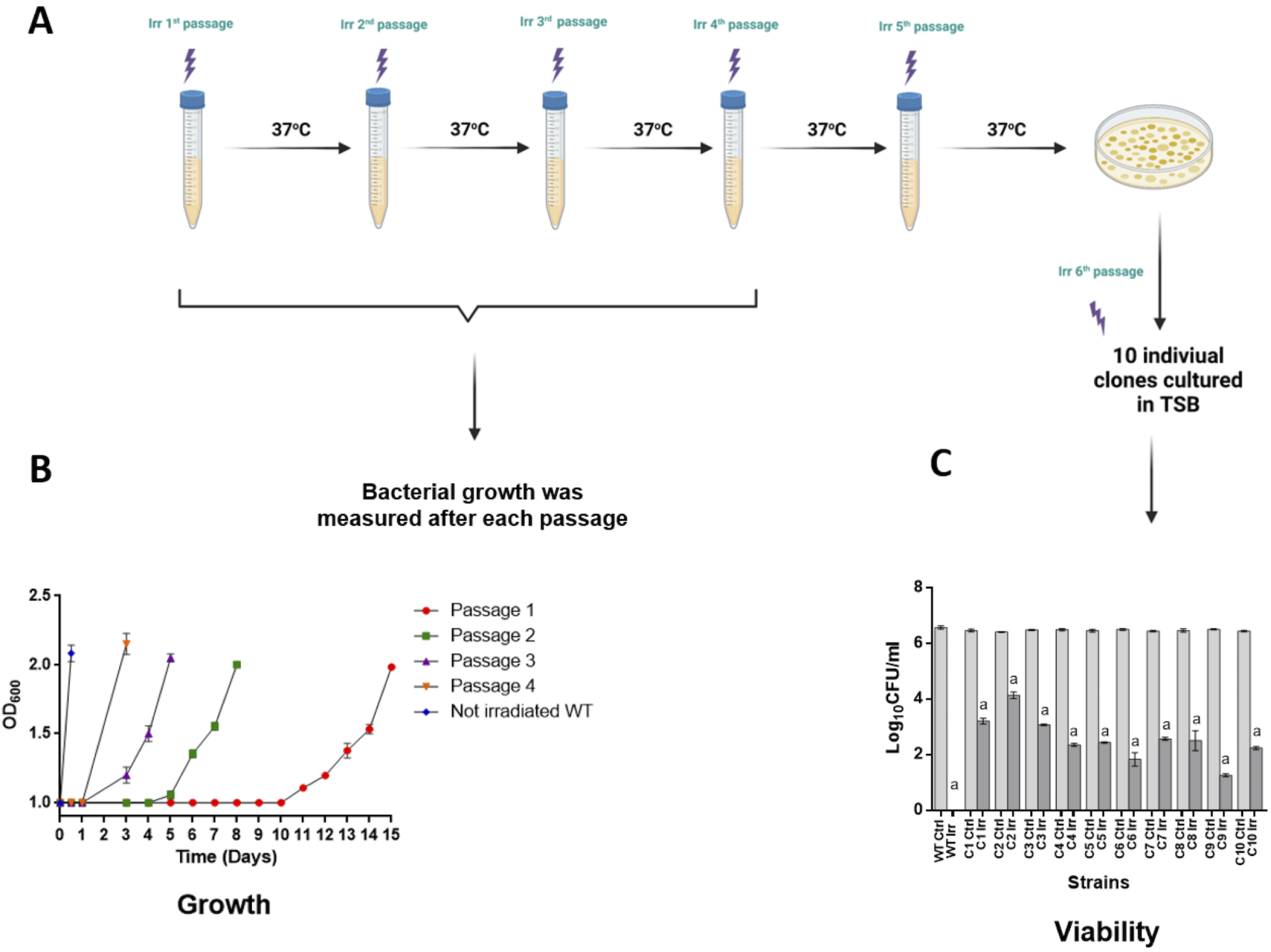
*E. coli* O157:H7 adaptation to an initially lethal dose of irradiation. **A**: Procedure for adapting bacteria to a lethal dose (1.5 KGy). Bacteria were adapted to1,5 kGy **γ*-***irradiation dose by successive passages from the exponential phase of growth (OD_600_ ≈ 1). After each passage, bacteria were incubated at 37°C with shaking until OD_600_=2. Bacterial cultures were subjected consecutively to a radiation dose of 1.5 kGy. After the 5 ^th^passage, bacterial cells (OD_600_ ≈ 1) were irradiated, plated on TSA plates and incubated overnight at 37°C. **B**: Bacterial growth following irradiation after each passage. 200 μL of overnight (O/N) cultures were diluted to an OD_600_ = 0.05 in TSB, were distributed into wells of sterile microtiter plates and incubated at 37°C for 20 h without agitation. The OD_600_ growth measurements were performed by Bioscreen C apparatus every hour after a mixing period of 30 seconds. **C**: Viability of ten adapted clones after the 6 ^th^ passage following irradiation. Clones culture were incubated individualy O/N at 37°C with shaking, diluted 100 time, incubated at 37°C with shaking until OD_600_ ≈ 1, irradiated at 1,5 kGy, serially diluted, plated on TSA plates and incubated overnight (O/N) at 37°C. The decrease in viability after irradiation, compared to the control was significant a=(P<0.001) for all tested samples. The results represent the means of replicate experiments for a minimum of three samples. Differences between the log_10_CFU counts of the Irradiated (Irr)/control untreated (Ctrl) for each strain vs wild-type (WT) parental strain were analyzed using Student’s t test.

Whole genome sequencing revealed that *E. coli* O157:H7 C1, C2, and C3 irradiation-adapted clones were found to have deletions of the two prophage, CP-933V and BP-933W (encoding the Stx1 and Stx2 toxins) DNA sequences and of two genes (*lexA*, and *dinI*) after the second passage of irradiation. Further, three other genes, *rpoA, wrbA* and *molY*, which encodes a hypothetical protein, contained point mutations (**Figure 2**). PCR amplification and sequencing of *rpoA, wrbA* and *molY* from sequential clones obtained after each passage showed that these mutations appeared after the second or third passage. For example, in *wrbA*, nucleotide G at position 25 was substituted by A (Valine at position 9 by Isoleucine) after the second passage (**Table 2**).

**Figure 2.**
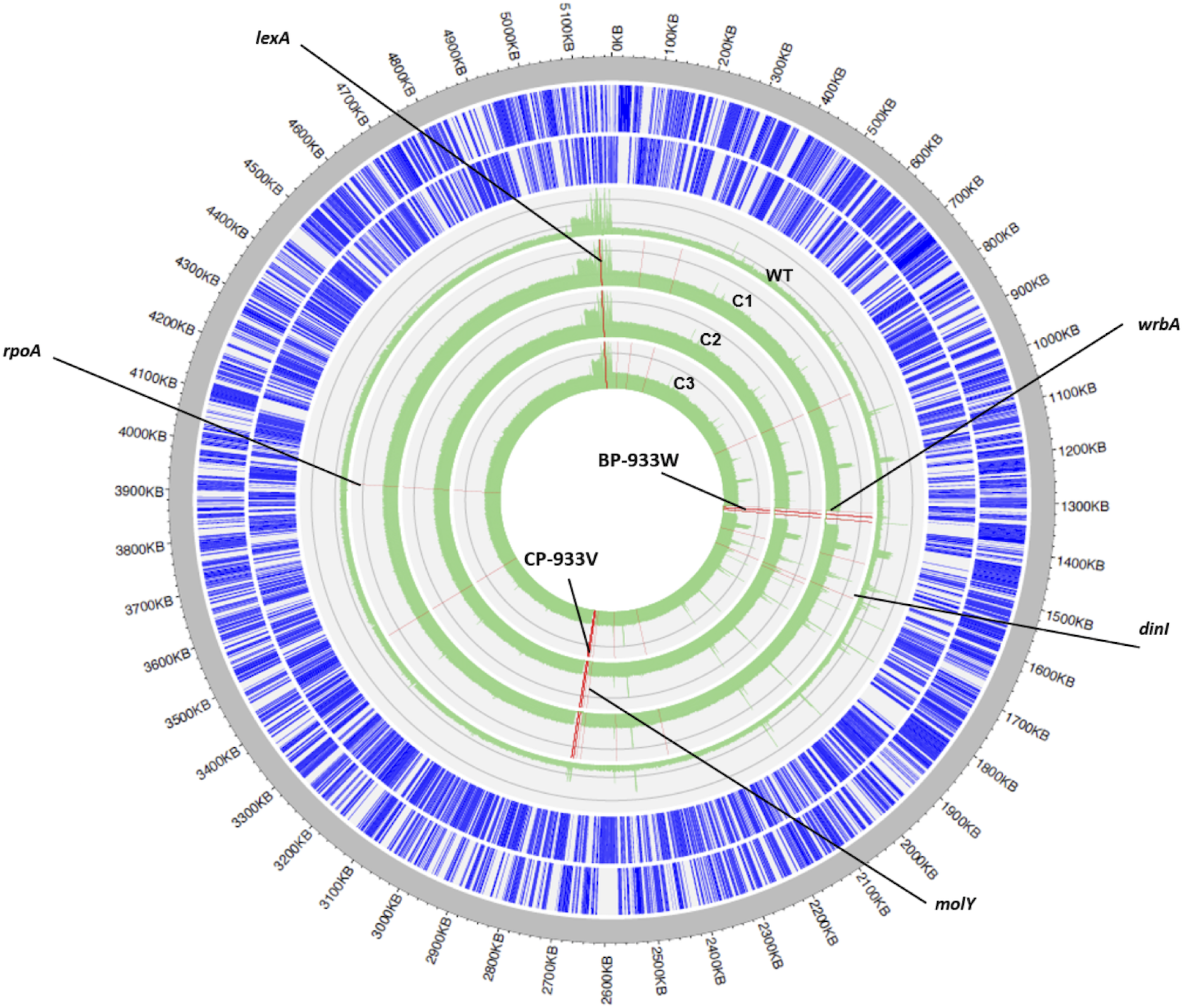
Circular genome maps of *E. coli* O157:H7 strain EDL933 and clones adapted to a lethal dose of irradiation (C1, C2 and C3). From outside to inside the wild-type followed by the C1, C2, and C3 clones respectively (green circles), the circles represent the genes encoded on the forward strand, those encoded on the reverse strand, the coverage of the wild-type and that for the clones C1, C2 and C3. Common mutated genes between C1, C2 and C3 are indicated by red lines.

### Correlation between irradiation adaptation and loss of *stx* toxin encoding genes

Sequencing data suggested that loss of the prophages could be a beneficial adaptation to survive irradiation stress. To investigate this hypothesis, *E. coli* O157:H7 strain Sakai (isolated from a human clinical case) and four other isolates of *E. coli* O157:H7 from animals or humans were also tested to validate the correlation between adaptation to a lethal dose of γ-irradiation and loss of *stx* genes. Confirmation of identification and serotype of the four isolates including 2 non-toxinogenic O157:H7 isolates was performed by Quebec Public Health laboratory using anti-O157 and H7 antisera. The strains were subjected to molecular typing by Pulsed Field Gel Electrophoresis according to procedures approved by Pulsenet international [24].

Isolates 2 and 3 are *stx*-free and could survive immediately following irradiation treatment at 1.5 kGy. The other strains lost their *stx* genes after the second passage of irradiation (**Table 3**).

### Lysogeny by prophage BP-933W and gain in cytotoxicity

To further confirm the hypothesis that a gain in resistance to irradiation was due to prophage excision, each of the three adapted clones and the *E. coli* K-12 strain MG1655 (as a control) were incubated with the culture supernatant of irradiated *E. coli* O157:H7 WT for 18h at 37°C to obtain lysogens that had been reinfected with prophages [25]. Lysogenic clones of C1, C2 and of *E. coli* K-12 MG1655 (C1-Φ, C2-Φ, and K-12-Φ respectively) were isolated and the presence of shiga toxin genes *stx*1 A/B, *stx*2 A/B, *lexA* and *dinI* in the whole DNA (genomic and circular) of these lysogens was investigated by PCR. We were unable to obtain a lysogen of clone C3. PCR demonstrated that the lysogens had gained *stx2* genes, as well as *lexA* and *dinI* genes. Genomic DNA sequencing revealed the absence of the phages in the C1-Φ, C2-Φ bacterial genomes. In contrast, BP-933W was integrated in E. coli K12-Φ at the *wrbA*-specific CATCGTTTCAATATGTC site. This site has previously been recognized for the integration of bacteriophage 933W, in *E. coli* K-12 [26, 27]. Interestingly, both *lexA* and *dinI* genes were also present in at the same positions as the EDL933 WT strain: *lexA* from 5140283 pb to 5140891 pb between *dgkA* and *dinF* genes; *dinI* from 1565472 pb to 1565717 pb between *yceP* and *pyrC* genes.

The production of Shiga toxins was evaluated by ELISA. As expected, C1 and C2 did not produce Shiga toxin. In contrast, these clones produced Shiga toxin after lysogenization by phage (C1-Φ and C2-Φ) (**Figure 3A**). The cytotoxicity effect of clones (C1, C2), and lysogens (C1-Φ and C2-Φ) vs *E. coli* O157:H7 WT was determined (**Figure 3B**). According to previous research, STEC strains are cytotoxic for Vero cells [28]. To evaluate this phenotype, Vero cells were exposed to supernatants of bacterial cultures as described in materials and methods. The lactate dehydrogenase (LDH) released from damaged Vero cells was measured, as described in the CytoTox96 kit (Promega, USA). The amount of LDH released is proportional to the number of lysed Vero cells. Strains producing Stx toxin gave higher cytotoxicity levels after 12-hours of incubation, However, cytotoxicity was completely absent when Vero cells were infected with clones C1 or C2. Interestingly, K-12-Φ was also able to produce Stx and had become cytotoxic. According to the sequence data, the prophage BP-933W was suspected to be present as an episome in *E. coli* C1-Φ and C2-Φ. To confirm this, extra-chromosomal DNA from *E. coli* C1-Φ and C2-Φ was extracted and analyzed by pulsed-field gel electrophoresis (data not shown). A low concentration (< 1ng/ul) of an extrachromosomal element of 61.7 KB was extracted from the gel. PCR of the extracted band confirmed detection of *stx2* (A&B) from the extracted extrachromosomal DNA. Cytotoxicity test results confirmed that the presence of Stx toxin could damage Vero cells, induce cell lysis, and release lactate dehydrogenase [29].

**Figure 3.**
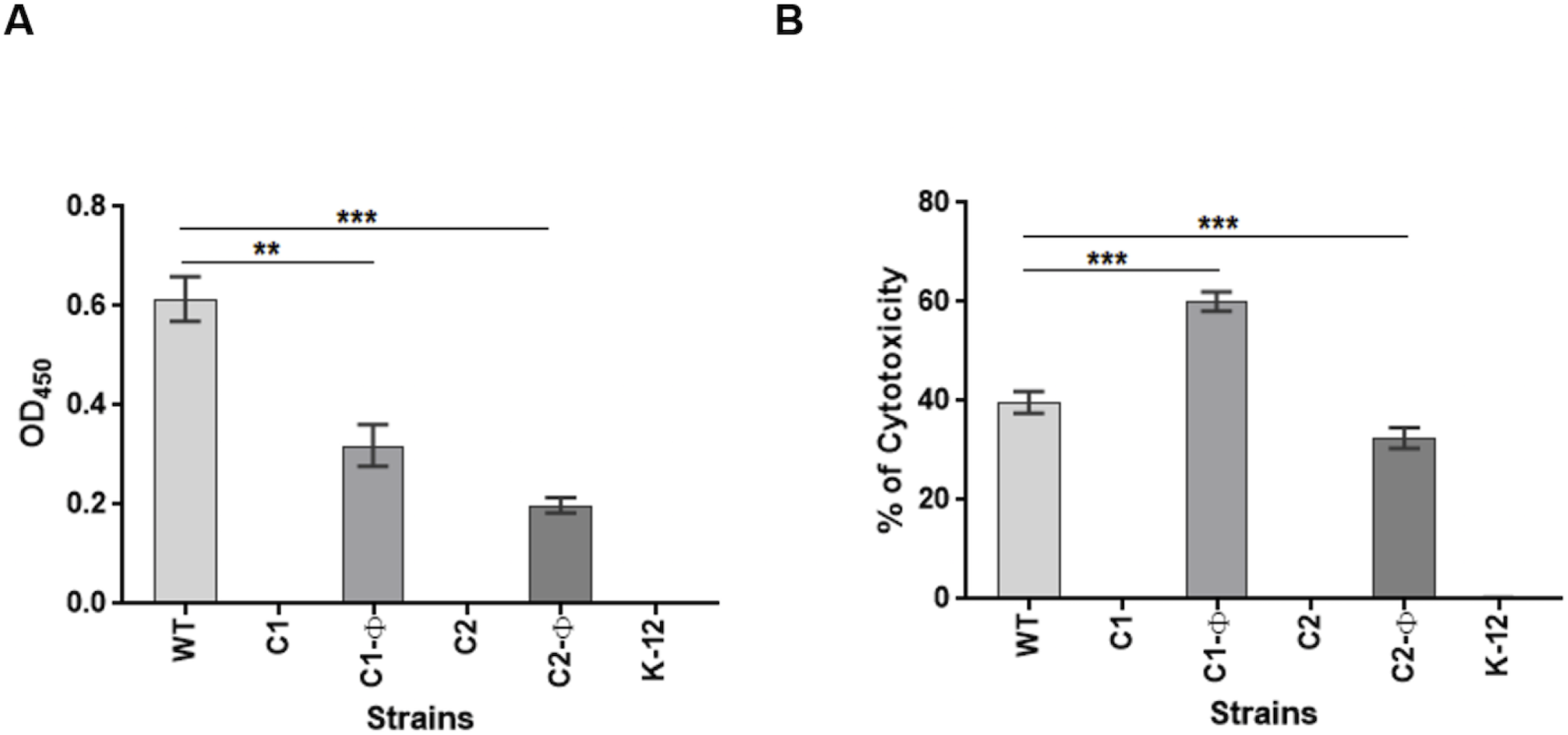
Cytotoxicity of *E. coli* O157:H7 clones carrying or not BP-933W prophages. **A**: Shiga Toxin detection in supernatants of *E. coli* O157:H7, C1, C2, K-12 MG1655, C1-Φ, C2-Φ and K-12-Φ by ELISA test. Each strain was incubated in TSB at 37°C (O/N) diluted 1/10 in TSB supplemented with mitomycin C (50 ng/mL) and was incubated at 37°C for 20 hours. ELISA was performed in 24-well microplates using 1 mL of each supernatant, Rabbit IgG, anti-Shiga Toxin 2 as detection antibody and polyclonal anti-rabbit as secondary antibody coupled to Horseradish peroxidase. Absorbance was read using a Biotek microplate reader at 450 nm and Gen 5 2.07 software. **B**: Cytotoxicity effect on VERO cells using LDH Release detection test. Vero cells were plated into 96-well cell culture plates in Dulbecco’s Modified Eagle’s medium supplemented with 10% heat-inactivated fetal bovine serum and incubated at 37°C under a 5% CO_2_ atmosphere. Cells were washed with Hanks’ Balanced Salt solution 1X and bacteria were added to confluent monolayers at a multiplicity of infection of 100 for 2, 4, 9 and 16 h. The total amount of LDH released into the medium was determined using the CytoTox96 kit. The percentage of cytotoxicity was calculated using the following formula: Percent cytotoxicity = 100 × Experimental LDH Release (OD490)/ Maximum LDH Release (OD490). The results represent the means of replicate experiments for a minimum of three samples. Differences between each strain vs WT were analyzed using Student’s t test (* p < 0.05, **p < 0.01, ***p < 0.001).

### Resistance to a lethal dose of irradiation

A dose of 1.5 kGy of irradiation is known to be lethal for *E. coli* O157:H7 [30]. At the same time clones C1, C2 and C3 adapted, and over time through six repeated passages and exposure to this irradiation dose, were able to become more resistant and grow following this high level of radiation stress (**Figures 1C and 5A**). Interestingly, *E. coli* K-12 strain MG1655 was more resistant to irradiation than wild-type *E. coli* O157:H7 strain EDL933 and showed a survival rate of 4.0 log_10_ CFU/mL (**Figures 4 and 5C**). Compared to *E. coli* O157:H7 EDL933, which showed no growth at 1.5 kGy of irradiation, the C1 clone survived with viable counts decreasing to 2.86 log_10_ CFU/mL and was able to grow after an adaptive period of 8 h. In contrast, prophage reintroduction reduced the viability and growth of C1 (**Figures 4 and 5B**). Immediately following irradiation treatment, the lysogen C1-Φ only demonstrated survival of 0.5 log_10_ CFU/mL (**Figure 4**). Similarly, prophage introduction increased sensitivity of *E. coli* K-12 to irradiation. Viable counts of K-12-Φ lysogen decreased to 0.98 log_10_ CFU/mL after exposure to 1.5kGy (**Figures 4 and 5C**).

**Figure 4.**
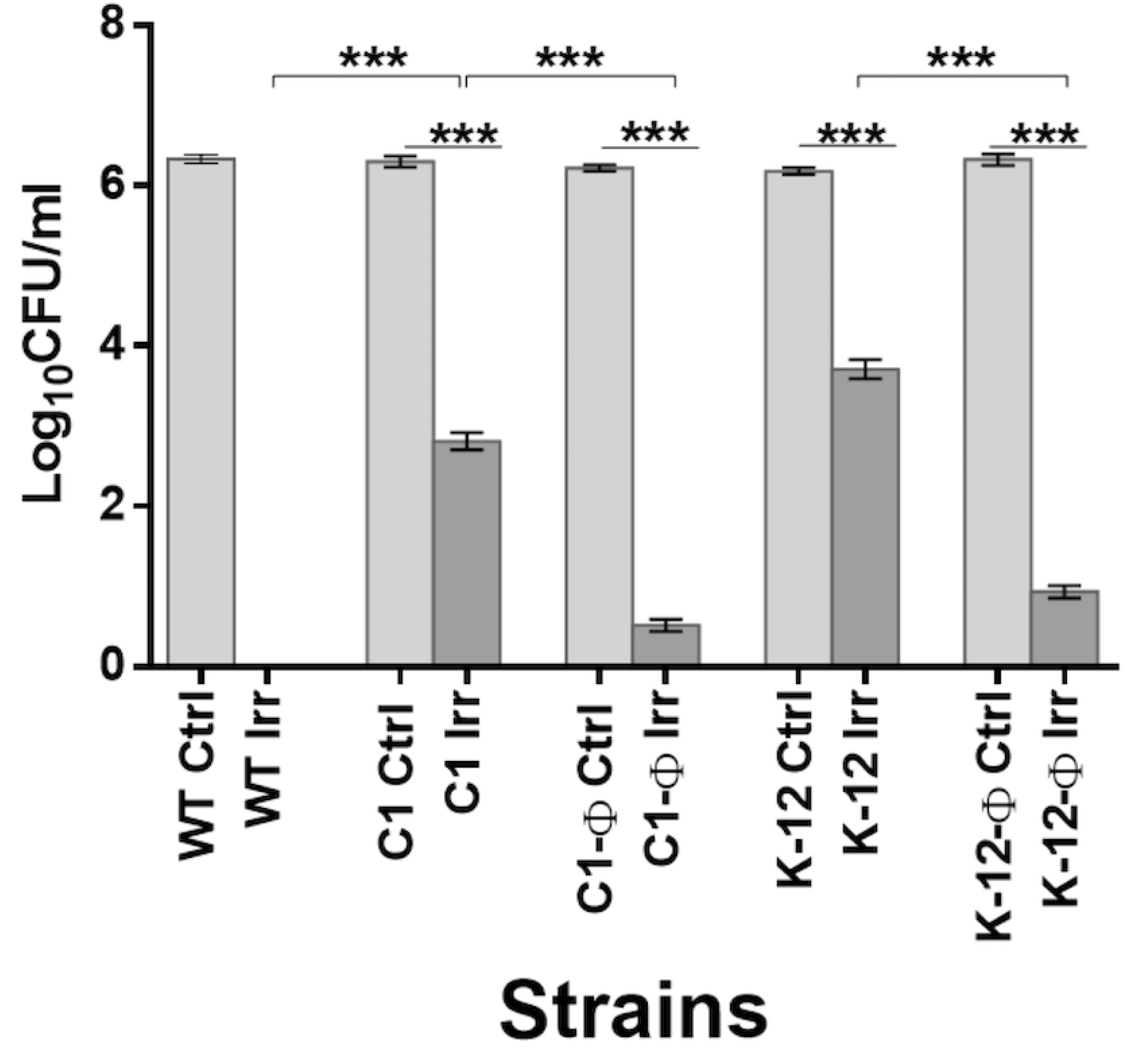
Viability of *E. coli* O157:H7 clones carrying or not BP-933W prophages following a lethal dose of γ-irradiation. Bacterial culture was incubated O/N at 37°C with shaking, diluted 100 times, incubated at 37°C with shaking until OD_600_ ≈ 1, irradiated at 1,5 kGy, serially diluted, plated on TSA plates and incubated overnight (O/N) at 37°C. The wild-type *E. coli* O157:H7 EDL933 showed no growth at 1.5 kGy of irradiation compared to the C1 that was able to survive and grow after an adaptive period of 8 h. However, the introduction of prophage reduced the viability and growth of C1-Φ. Similarly, prophage introduction increased sensitivity of *E. coli* K-12 to irradiation after exposure to 1.5kGy. Ctrl: control non-irradiated; Irr: irradiated. The results represent the means of replicate experiments for a minimum of three samples. Differences between the log_10_CFU counts of (Irr /Clrt) for each strain were analyzed using Student’s t test (* p < 0.05, **p < 0.01, ***p < 0.001).

**Figure 5.**
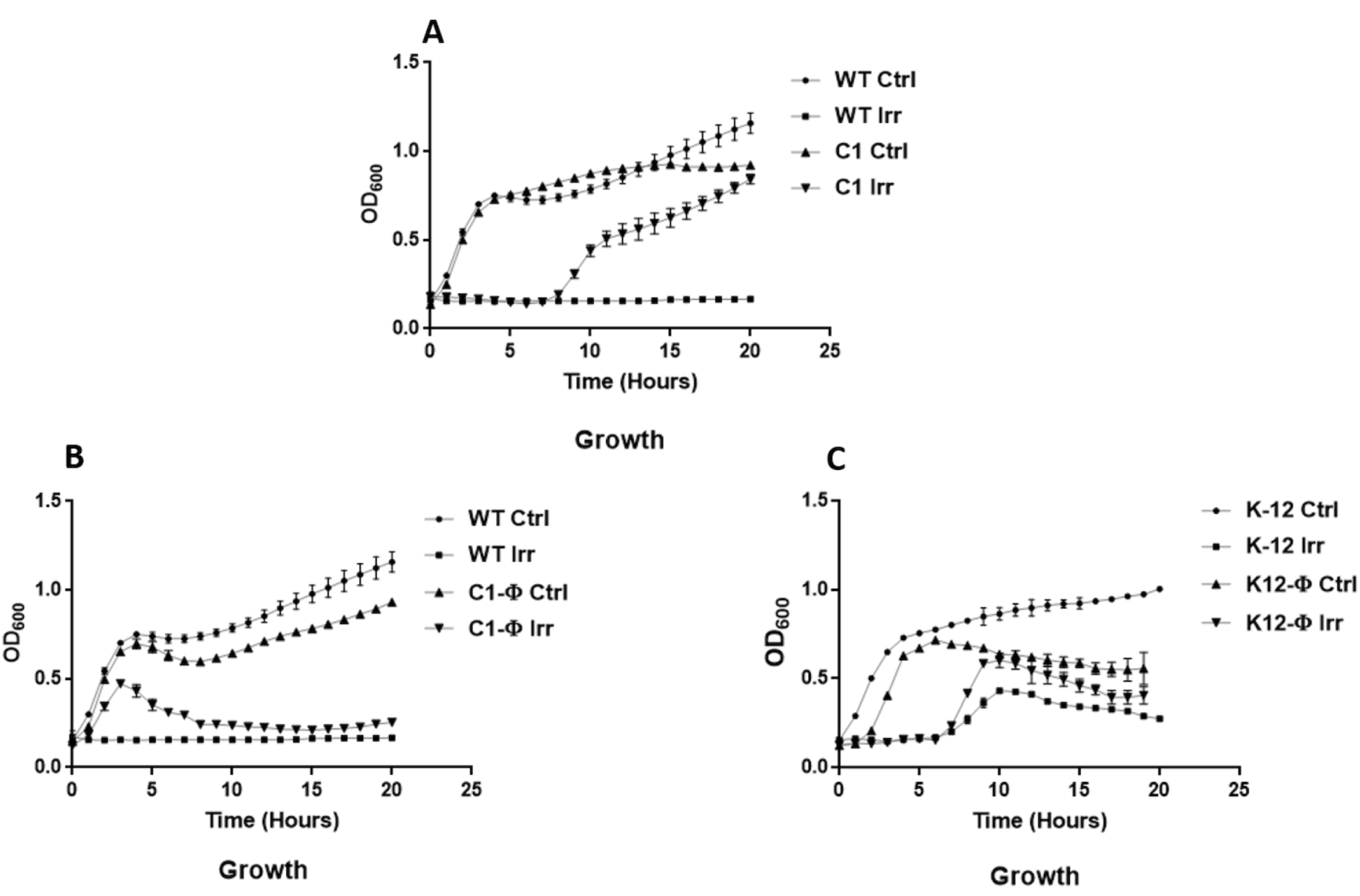
Growth of *E. coli* O157:H7 adapted to a lethal dose of irradiation. Growth of (A) C1, (B) C1-Φ or (C) K-12-Φ. O/N bacterial culture was diluted 100 times, incubated at 37°C with shaking until OD_600_ ≈ 1, irradiated at 1,5 kGy. 200 μL of cell cultures diluted to an OD_600_ = 0.05 in TSB, were distributed into wells of sterile microtiter plates and incubated at 37°C for 20 h without agitation. The OD_600_ growth measurements were performed by Bioscreen C apparatus every hour after a mixing period of 30 seconds. The results represent the means of replicate experiments for a minimum of three samples. Ctrl: non-irradiated control; Irr: irradiated. Vertical bars represent the standard errors of the means. Statistical significance was calculated by Student’s t test.

### Acid resistance

A hallmark of the pathogenicity of *E. coli* O157:H7 is its resistance to acid in the gastrointestinal tract [31]. To verify if clones that lost the PV-933V and BP-933W prophages during adaptation to irradiation become more sensitive to acidic pH, we determined bacterial viability in TSB medium at pH=2. *E. coli* O157:H7 radioresistant clone C1 did not survive after 2h in this acid pH environment, whereas the parental strain EDL933 was resistant to acid (pH=2), as expected. Interestingly, prophage re-introduction increased significantly the viability (P≤0.005) up to 4 h for the C1-Φ lysogen (**Figure 6**). However, the viability of C1 after re-introduction of prophages was still less than viability of the EDL933 parent strain when exposed to acidic pH 2 environment. Introduction of phage BP-933W to *E. coli* K-12 also increased survival at pH 2 particularly after 3 to 4 hours of treatment. These results suggest that bacteriophages and particularly BP-933W may contribute to increased acid resistance in *E. coli* O157:H7 EDL933 and *E. coli* K-12 strains.

**Figure 6.**
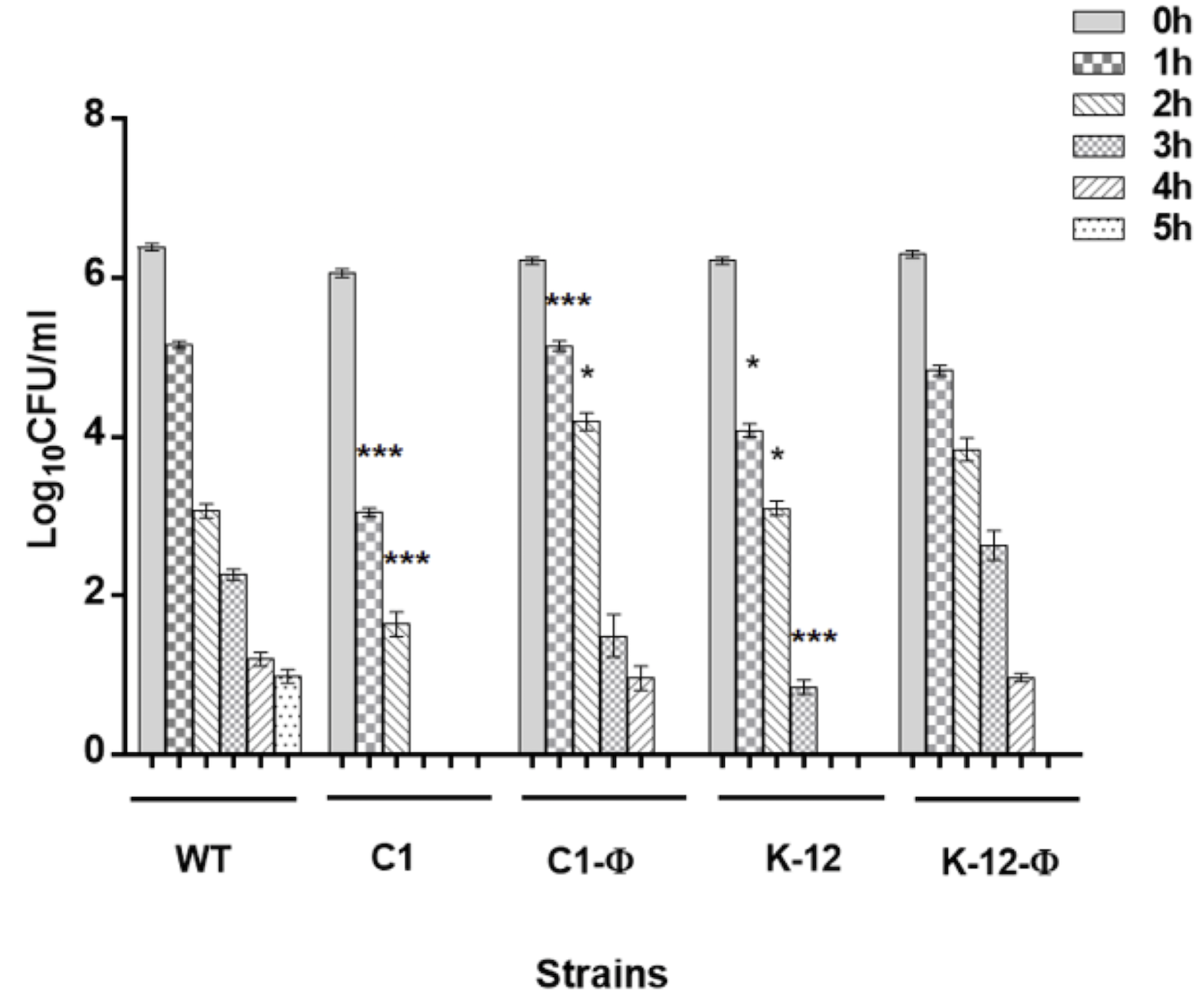
Viability of *E. coli* O157:H7 clones carrying or not BP-933W prophages and subjected to acid stress (pH 2) over time. O/N bacterial culture was diluted to an OD_600_ ≈ 1 in TSB at pH=2 (HCl, 1N) and incubated at 37°C under agitation for 5 h. Bacteria were serially diluted every hour, plated on TSA plates and incubated overnight (O/N) at 37°C. The results represent the means of replicate experiments for a minimum of three samples. The log_10_CFU counts for each strain vs WT were analyzed every hour using Student’s t test. There is no significant difference between each strain (C1, C-phage) and WT and between K-12 and K-12-phage at the beginning of the test (* p < 0.05, **p < 0.01, ***p < 0.001).

### Paraquat resistance

As with irradiation stress, oxidative stress produces reactive oxygen species (ROS) which can cause DNA damage and activate a bacterial stress response [32] and also induce activation of the bacteriophage lytic cycle [33]. To determine if prophages contribute to aggravating an ROS response, we tested paraquat resistance of *E. coli* O157:H7, the clones that were adapted to irradiation and clones wherein prophages were re-introduced. As shown in **Figure 7**, parental strain EDL933 is sensitive to paraquat and its viability was reduced by 0.3-1.4 log_10_ CFU/mL. By contrast, radioresistant clones C1 and C2 were completely resistant to paraquat stress, indicating a correlation between oxidative and irradiation stress resistance. Further, re-introduction of prophage in these radiation-adapted clones (lysogenic strains C1-Φ and C2-Φ) rendered them significantly sensitive to paraquat (P≤0.05 with 0.2 µM and P≤0.005 with 0.3 and 0.5 µM) compared to the adapted clones C1 and C2. Viability of these bacteria was reduced by 1.4-1.8 log_10_ CFU/mL after exposure to paraquat. Introduction of prophage BP-933W to *E. coli* K-12 also resulted in an increased sensitivity to paraquat at 0.2 µM, wherein *E. coli* K-12-Φ had a 1 log greater decrease in viability when compared to parental strain *E. coli* K-12 strain MG1655. These results indicate that the presence of bacteriophage, such as shiga toxin encoding phages, in addition to rendering *E. coli* more sensitive to radiation stress, can also increase sensitivity to oxidative stress.

**Figure 7.**
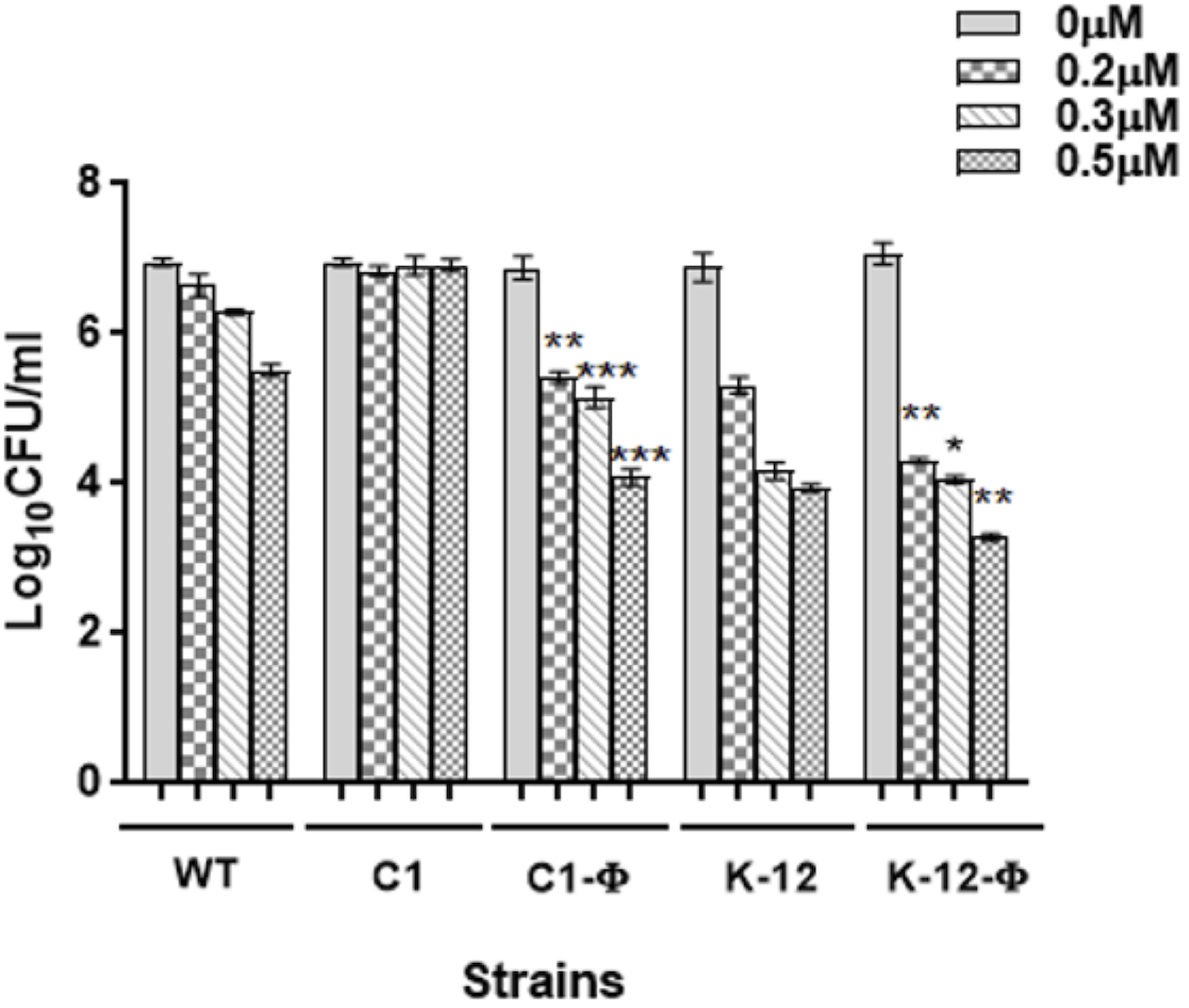
Viability of *E. coli* O157:H7 clones carrying or not BP-933W prophages under oxidative stress (paraquat). O/N bacterial culture was diluted to an OD_600_ ≈ 1 was incubated in TSB supplemented with paraquat at 0.2µM, 0.3µM or 0.5µM, incubated at 37°C under agitation for 40 min, serially diluted, plated on TSA plates and incubated overnight (O/N) at 37°C. The log_10_CFU counts for each strain vs WT were analyzed using Student’s t test. The results represent the means of replicate experiments for a minimum of three samples. There is no significant difference between each strain (C1, C-phage) and WT and between K-12 and K-12-phage at the beginning of the test (* p < 0.05, **p < 0.01, ***p < 0.001).

## Discussion

Irradiation can effectively reduce or eliminate some bacterial pathogens from foods, although some bacteria demonstrate an adaptive response and resistance to radiation treatment. How EHEC can adapt to normally lethal doses of γ-irradiation is not well understood. In addition, one of the key virulence traits of EHEC O157:H7 is the production of Shiga toxins (Stx1 and Stx2) encoded respectively by CP-933V and BP-933W temperate bacteriophages [7, 9, 34]. Oxidative stress has also been shown induce shiga toxin encoding lambdoid phages and as such EHEC lysogens harboring these phages can be more sensitive to oxidative stress [33, 35]. In this work, the potential for adaptation of O157:H7 strain EDL933 to an initially lethal dose of irradiation was investigated by repeatedly irradiating bacterial cells followed by a period of growth restoration over six cycles. It was found that selected radioresistant clones also had increased resistance to paraquat-generated oxidative stress and were more sensitive to acidity, compared to parental strain EDL933. Loss of both BP-933W and CP-933V prophages (which also resulted in loss of the *stx1* and *stx2* toxin encoding genes) was responsible for the adaptation to a lethal dose of irradiation in these clones. This result is coherent with reports describing the loss of the CP-933V and BP-933W *E. coli* O157:H7 prophages from the genome following antibiotic stress from exposure to ciprofloxacin, fosfomycin, or mitomycin C, although this stress also promoted bacterial cell lysis [36].

There is increasing evidence that Stx prophage-encoded factors can modify bacterial gene expression and alter multiple phenotypes. Shiga toxin phage lysogeny has a direct effect on the global expression of bacterial genes; moreover, an increase in acid tolerance and motility has been reported when bacteria were lysogenized [37]. Further, RNA sequencing (RNA-seq) studies have revealed a positive effect of the phage ϕ24B carrying *stx2a* on the gene expression of acid resistance in *E. coli* K-12 strain MC1061 mediated by the phage-encoded regulator CII. Moreover, CII also apparently represses expression of elements of the LEE-encoded type III secretion system, which is critical for EHEC virulence [38, 39].

Several studies have shown that phage integration or excision can inactivate or alter gene function through physical disruption of the genes or their promoter regions. As such, insertion and excision of prophages from bacterial genomes can alter bacterial physiology [40]. CP-933V is integrated near the *mlrA* (*yehV*) chromosomal locus and contributes to reduced expression of curli fimbriae and biofilm formation [41]. BP-933W is integrated at the *wrbA* site and alters responses to environmental stress [42, 43]. Deletion of these prophages may therefore lead to changes in *mlrA* and *wrbA* expression and may restore their native function and regulation in *E. coli* [44].

In our study, CP-933V and BP-933W excision was an early adaptation to survive γ-irradiation stress, as it occurred after the second irradiation cycle. Under DNA-damage stress, RecA cleaves the CI repressor promoting phage induction which can lead to phage release and bacterial cell lysis [45]. Some bacteria may excise the prophages as an episome that may potentially reintegrate into the bacterial genome. For certain phages, the episome can also be lost without killing the bacteria [40]. This is the case with phages CP-933V and BP-933W which were excised and were eliminated from clones (C1 and C2) after 2 passages at 1.5kGy. In our current study, it is likely that the increase in resistance of the adapted clones to γ-irradiation and oxidative stress was due to loss of the prophage genomes by excision from the bacterial chromosome. In addition, the deletion of *lexA* and *dinI*, which constitutive activation of the SOS response, also occurred when phage genomes were lost.

Interestingly, in the process of adaptation of *E. coli* K-12 to a lethal dose of irradiation, among other genomic alterations, the *E. coli* K-12 prophage e14 was also deleted from the genome [46, 47]. As with CP-933V and BP-933W, e14 is a lambdoid prophage that is inducible by mitomycin C treatment. The insertion site in the bacterial genome is the isocitrate dehydrogenase (*icd*) gene that is involved in oxidative stress resistance [36, 47, 48]. These data corroborate our results and indicate that increased resistance to irradiation in *E. coli* is linked to a loss of prophage elements from the genome.

In addition, the e14 prophage is excised in the response to SOS induction [49]. Harris *et al*. [49] have concluded that this phenomenon is due to mutations in the *recA* gene. In the present work, *E. coli* O157:H7 adapted to a lethal dose of γ-irradiation by losing the *lexA* and *dinI* genes. However, no mutations were identified in the *recA* gene in radiation-adapted clones C1, C2 and C3 (figure S1). The possible explanation for CP-933V and BP-933W excision is likely an increased promotion of the SOS response via *lexA* and *dinI* deletion which can promote consecutive expression of *recA* after irradiation exposure.

It is possible that *E. coli* O157:H7 radiosensitivity may therefore in part be due to prophage induction, which could result in increased bacterial cell death and lysis. To test this hypothesis CP-933V and BP-933W prophages were re-introduced into *E. coli* O157:H7 radioresistant clones C1 and C2 using the method of Hull, Acheson [50]. It was shown that only BP-933W could be re-introduced into the bacterial clones (C1 and C2), but the phage did not re-integrate back into the bacterial genome. Moreover, to re-introduce prophage, clones and strains were incubated with the supernatant of irradiated *E. coli* O157:H7 WT. It was, however, not determined whether this supernatant contained both CP-933V and BP-933W or only BP-933W phages. Interestingly, re-introduction of BP-933W to the radiation-adapted clones C1 and C2 was accompanied by a regain of the *lexA* and *dinI* genes. *E. coli* O157:H7 C1, C2 and C3 adapted clones, wherein *lexA* and *dinI* genes are deleted, and are radioresistant compared to *E. coli* O157:H7 parental strain EDL933. Although, the relation between the combined loss of both the prophages and these two genes from the adapted clones remains to be identified, it might be that *lexA* and *dinI* are required for re-introduction of BP-933W into the bacteria, the genom. since LexA is a key regulator of the SOS response and regulation of the phage temperate/lytic cycle in *E. coli* and other bacteria [51] and DinI inhibits RecA, and suppresses the SOS response [52].

It is important to highlight that, as observed in clones C1, C2 and C3, two passages at a 1.5 kGy lethal dose of irradiation also caused the loss of *lexA, dinI* and *stx* genes (1 and 2) in other EHEC strains including isolate 1, isolate 4 and the EHEC Sakai strain (**Table 3**). de Vargas and Landy [53] have shown that simultaneous excision of the lambdoid prophages inhibits reintegration of phages by disruption of the phage attachment site (*attP*). In this study, the bacterial *attB* site in the *wrbA* gene [42] is mutated (G25 >A) suggesting that mutation of the bacterial attachment site may induce prophage excision and inhibit further integration at the same site. Also, other studies [54, 55] showed for other Stx-phages an ability to integrate into multiple sites in a host bacterial strain that may be even sometimes resolved in polylysogeny and increased Stx-toxin production. For example, for the stx-phage Φ24_B_ that shares a considerable amount of genomic DNA sequence with Stx-phage 933W several alternate integration sites were identified when the *attB* site in the *wrbA* gene was altered or deleted. Most of these alternative sites are located in genes encoding proteins of unknown function. So, since *wrbA* was mutated following irradiation and passaging, this may explain why BP-933W could not integrate at the altered *wrbA* attachment site.

Notably, the clones which have been adapted to radiation treatment have lost the shiga toxin encoding genes and are now no longer cytotoxic to Vero epithelial cells. The cytotoxicity, however, is restored following reintroduction of BP-933W and expression of the Stx2 toxin. Although clone C1 is non-cytotoxic, it is resistant to a lethal dose of irradiation. By contrast, the C1-Φ lysogen that regained BP-933W and *stx2* significantly lost radioresistance concomitant with a regain in the capacity to kill target host epithelial cells. Taken together, this indicates that while harboring the *stx2* encoding prophage, can contribute to EHEC virulence/host cell toxicity, its carriage is detrimental for surviving radiation stress. Taken together, results support the use of γ-radiation to effectively reduce and eliminate EHEC from food products, but demonstrate a low, but potential capacity for such strains to develop radio-resistance. Importantly however, the potential adaptation of EHEC to high doses of radiation would result in elimination of the Stx encoding phages and would likely lead to a decreased resistance to acid pH and would greatly reduce the potential of such strains to cause disease.

## Conclusion

In this work, *E. coli* O157:H7 adaptation to a typically lethal dose of γ-irradiation involved modifications in the genome including deletion of CP-933V and BP-933W prophages encoding the Stx toxins. Further, loss of these prophages resulted in loss of cytotoxicity to epithelial cells and significantly decreased resistance to acidity (pH 2.0) while increasing resistance to oxidative stress. This specific adaptation to irradiation might be seen as an example of a biological trade-off where the bacterium cannot maintain virulence and resistance at the same time. Accordingly, re-integration of prophage BP-933W in radioresistant clones resulted in both a regain in cytotoxicity and acid resistance, but a marked increase in radiosensitivity. Overall, results demonstrate that although there is some potential for EHEC to develop radioresistance, the adaptation would likely engender a loss of key virulence traits such as Stx-mediated cytotoxicity and decreased resistance to acid pH, both traits that are paramount for the low infectious dose and cell and tissue damage that are a hallmark of EHEC pathogenesis.

## Acknowledgements

We thank the Natural Sciences and Engineering Research Council of Canada (NSERC) discovery grant program for its financial support, through grants RGPIN-2017-05947 to ML, RGPIN-2019-06642 to CMD, and RGPIN-2016-04940 to FJV. This work was also supported by grant MOP-142466 from the Canadian Institutes of Health Research (CIHR) to ED. FJV is a research scholar of the Fonds de Recherche du Québec – Santé. ED holds the Canada Research Chair in Sociomicrobiology. ATV received a Postdoctoral Fellowship from the NSERC. Kateryna Krylova was funded by a MITACS Elevation Postdoctoral Fellowship. We thank Nordion Int. Inc. for performing the irradiation treatment. The authors also acknowledge the staff of the Centre d’Innovation Génome Québec and Université McGill for the preparation of libraries and sequencing service.

## Materials and methods

### Bacterial strains and culture conditions

In this study, *E. coli* O157:H7 EDL933 and Sakai, K-12 MG1655 strains (our laboratory stock), and four distinct isolates of *E. coli* O157:H7 (previously isolated from animals or humans and provided to us by the Quebec Public Health Laboratory) were used. All bacterial cultures were grown in Tryptic Soy Broth (TSB, Difco, Becton Dickinson, Sparks, MD, USA) with or without agitation (240 rpm) in a TC-7 roller drum (New Brunswick) or on Tryptic Soy Agar plates (TSA, Difco, Becton Dickinson, Sparks, MD, USA) at 37°C. A Bioscreen C apparatus (Growth Curves USA) was used for growth assays. Briefly, 200 μL of bacterial cell cultures diluted to an OD_600_ = 0.05 in TSB, were distributed into wells of sterile microtiter plates and then incubated at 37°C for 20 h without agitation. The OD_600_ growth measurements were performed every hour after a mixing period of 30 seconds. For viability studies, bacterial cultures were serially diluted, plated on TSA plates and incubated overnight (O/N) at 37°C. For Shiga Toxin detection and cytotoxicity assays, 50 ng/mL mitomycin C (Sigma Aldrich, Ontario, Canada) was added to bacterial cultures.

### Adaptation of *E. coli* O157:H7 to a normally lethal dose of γ*-*irradiation

*E. coli* O157:H7 strain EDL933 was adapted to a normally lethal **γ*-*** irradiation dose by successive passages from the exponential phase of growth (OD_600_ ≈ 1). After each passage, bacteria were incubated at 37°C with shaking until OD_600_=2. Bacterial cultures were transferred into microfuge tubes and subjected consecutively to a radiation dose of 1.5 kGy [30]. After the 5^th^ passage, bacterial cells (OD_600_ ≈ 1) were irradiated, plated on TSA plates and then incubated overnight at 37°C. Ten colonies (clones) were cultivated O/N in TSB at 37°C, diluted 100 times, incubated at 37°C under agitation until OD_600_ ≈ 1 and irradiated at 1.5 kGy. Afterwards, viability and growth studies were performed. The irradiation treatments were performed at room temperature (20 ± 1 °C) at the Canadian Irradiation Center as described previously [56].

### Acid and oxidative stress resistance assays

*E. coli* O157:H7 wild-type strains, *E. coli* K-12 MG1655 and all clones with/without prophages were cultivated in TSB and incubated O/N at 37°C under agitation. For acid resistance assays, bacteria were diluted to an OD_600_ ≈ 1 in TSB at pH=2 (HCl, 1N) and incubated at 37°C under agitation for 5 h. Viability of bacteria was tested every hour as described for general culture conditions. For oxidative stress resistance, bacteria at OD_600_ ≈ 1 were incubated in TSB supplemented with paraquat at 0.2µM, 0.3µM or 0.5µM and incubated at 37°C under agitation for 40 min. Viability was then evaluated by plate count dilutions.

### Polymerase chain reaction (PCR)

Overnight bacterial cultures were diluted at 1/1000 and incubated in TSB at 37 °C until the exponential phase of growth (OD_600_ ≈ 1). DNA extraction was performed using DNeasy Blood and tissue Kit (Qiagen, Montreal, Canada) as recommended by the supplier. Amplification of genes was carried out by PCR using the specific primers that were designed using the Primer3 plus program (Table 1).

**Table 1.**
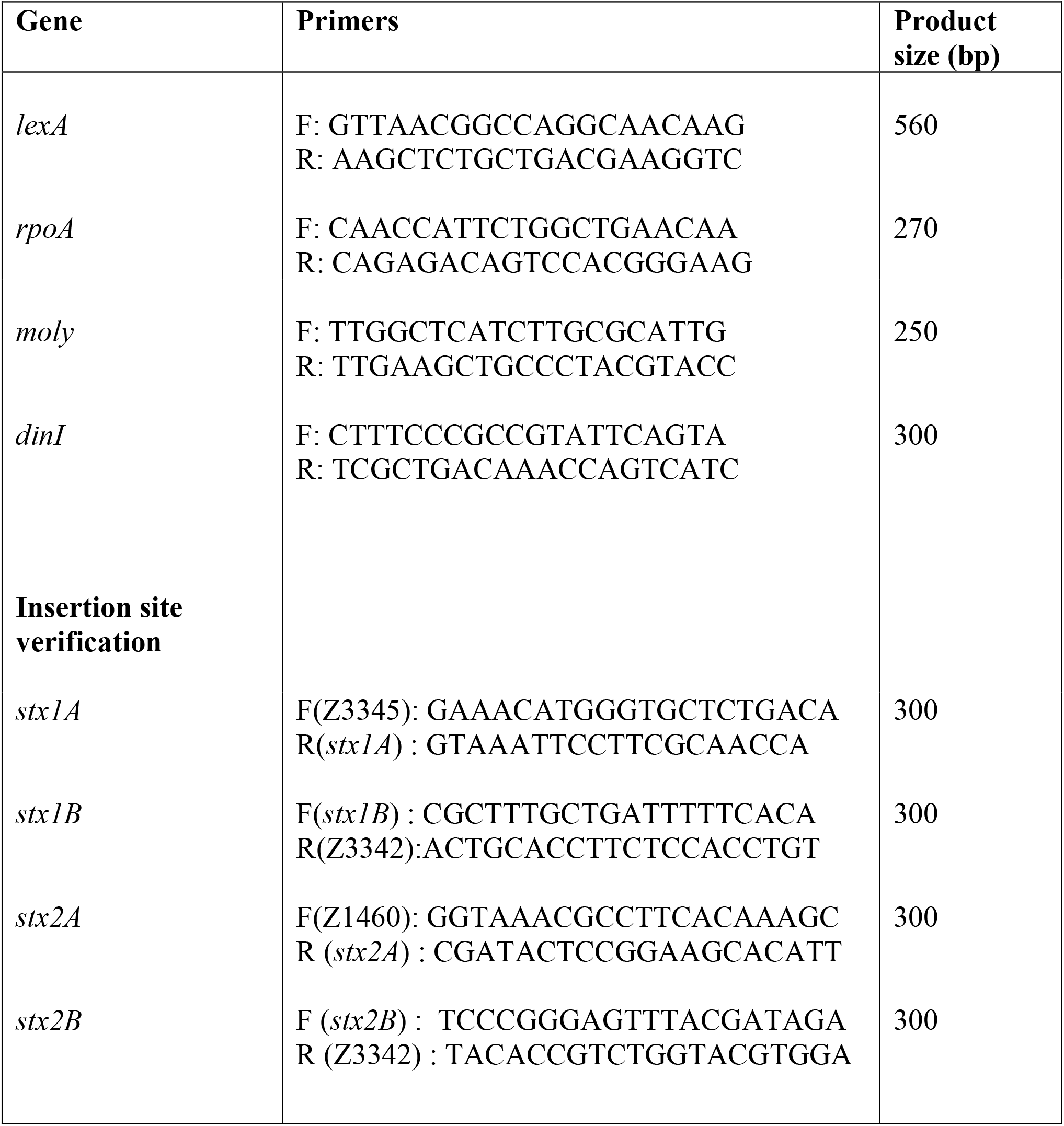
Primers used in this study.

**Table 2.** Point mutations identified in *E. coli* O157:H7 C1, C2 and C3 radiation adapted clones.

| Gene | Mutations | Passage |
| --- | --- | --- |
| **Wt_00837 (*wrbA*)** | p.G25A ; **p. Val9Ile** | P2 |
| **Wt_02639 (*molY*)** | p. C3109T ; **p.Arg1037Cys** | P3/P4 |
| **Wt_03821 (*rpoA*)** | p. G835T ; **p.Gly279Cys** | P1/P2 |

**Table 3.** Correlation between passage at the lethal dose of γ-irradiation and Stx carriage of *E. coli* O157:H7 strains.

| **Isolates or strains**<br>**(our lab)** | **Presence of *stx***<br>**before**<br>**irradiation** | **Time before**<br>**growth following**<br>**irradiation**<br>**(days)** | **Presence of *stx***<br>**After irradiation**<br>**(After passage 2)** |
| --- | --- | --- | --- |
| **Isolate 1 (Bovine)** | *stx1* and *stx2* | 8 | No |
| **Isolate 2 (human**<br>**case)** | None | 1 | No |
| **Isolate 3 (Bovine)** | None | 1 | No |
| **Isolate 4 (Pork)** | *stx1* and *stx2* | 21 | No |
| **Sakai (human case)** | *stx1* and *stx2* | 18 | No |

### Bioinformatics analyses

The DNA of the EHEC EDL933 wild-type strain, each of the adapted clones, and *E. coli* K-12 and lysogens of clones was sequenced at high-throughput using an Illumina HiSeq at the Sequencing Platform of Genome Canada (McGill University, Canada). The whole genome shotgun sequencing data set was deposited to the Sequence Read Archive of the National Center for Biotechnology Information under the BioProject PRJNA777472. The raw sequencing reads were processed using fastp version 0.19.5 [57] and ParDRe 2.2.5 [58]. Read quality was assessed using FastQC version 0.11.8 (https://www.bioinformatics.babraham.ac.uk/projects/fastqc/). The reads were *de novo* assembled using SPAdes version 3.13.0 [59] with k-mer lengths of 21, 33, 55 and 77. The resulting contigs were aligned on the reference sequence of *E. coli* EDL933 [60] using Mauve version 2015-02-13 [61] and were annotated using Prokka version 1.13.3 [62]. The nucleotide differences and genome alterations among adapted clones and wild-type strains were identified using Snippy version 3.2 (https://github.com/tseemann/snippy). The macro differences were visualized by mapping the reads from the clones onto the sequence of the wild-type strain with the combined use of BWA version 0.7.17-r1188 [63], SAMtools version 1.9 [64], and shinyCircos [65].

Statistical analyses were done using Prism 5 (GraphPad Software, Inc.). A *P* value was considered to be statistically significant when *P* ≤ 0.05 (for either analyses using the Student *t*-test or Kruskal-Wallis rank sum test).

### Cytotoxicity assays

Cytotoxicity assays were performed as described by Xiong *et al*. [28], with some modifications. Vero cells (ATCC^®^CCL-81™) were plated into 96-well cell culture plates (10^4^ cells/well) in Dulbecco’s Modified Eagle’s medium (Wisent Bioproducts, Canada) supplemented with 10% heat-inactivated fetal bovine serum (Wisent Bioproducts, Canada) and incubated at 37°C under a 5% CO_2_ atmosphere. Before infection, the cells were washed with Hanks’ Balanced Salt solution 1X (Wisent Bioproducts, Canada) and bacteria were added to confluent monolayers at a multiplicity of infection (MOI) of 100 for 2, 4, 9 and 16 h. The total amount of LDH released into the medium was determined using the CytoTox96 kit (Promega, USA) according to the manufacturer’s instructions and measured using a Tecan plate reader (Tecan Group Ltd) at 490 nm. The percentage of cytotoxicity was calculated using the following formula:

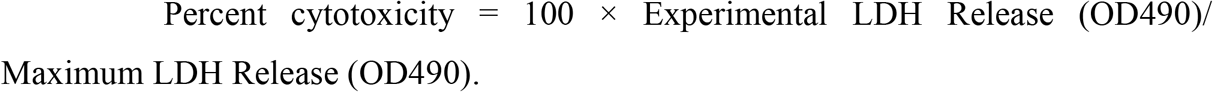

### Prophage re-introduction

*E. coli* O157:H7 EDL933 was cultivated in TSB supplemented with 50 ng/mL mitomycin C and incubated at 37°C for 18 h. Bacteria were centrifuged (5,000 x *g* for 10 min at 4°C) and the supernatant was filtered through syringe filters (0.45 μm pore size). Infection of bacteria with phages from the filtered supernatant was carried out as described by Lu and Breidt [25]. A quantity of 200 μl of TSB and 5 μl of *E. coli* K-12 MG1655 or of EHEC bacteria adapted to a lethal dose of γ-irradiation (Clone1: C1 Clone2: C2 Clone3: C3) at the exponential phase of growth (OD_600_ ≈ 1) and 45 μl of filtered supernatant were added and incubated in a 96-well microplate at 37°C for 20 h without agitation. To select lysogenic bacteria, 100 μl of cell culture from each well were centrifuged as described previously and 10 μl of each supernatant was used in spot-tests onto a lawn of *E. coli* K-12 MG1655 and by PCR for detection of *stx1*A/B, *stx2*A/B and *lexA* sequences using specific primers (Table 1).

### Pulsed-field gel electrophoresis (PFGE) of phage DNA

Phage DNA was extracted from O/N cultures of *E. coli* K-12 MG1655-Φ using the ZymoPURE Miniprep kit (Cedarlane, Burlington, Ontario, Canada). PFGE was performed with a CHEF Mapper XA from Bio-Rad with the following parameters: separation from 5 kb to 500 kb, calibration factor 1.0, buffer 0,5X TBE, temperature 14°C, 1.0% PFC Agarose, gradient 6.0 V/cm, run time 15 h 16 min, included angle 120°, Initial switch time 0.22s, final switch time 8.53 s, ramping factor Linear.

### Spot test

A 5 ml volume of targeted bacteria at OD_600_=0.15-0.2 was applied to TSA plates containing 0.5% agar for 30 sec and the culture was then discarded. A quantity of 10 µl of tested samples was spotted on the completely dried agar as a bacterial overlay. The plates were left to dry and were inspected for lysis zones after an O/N incubation at 37°C.

### Detection of Shiga toxins

Shiga toxins were detected in supernatants of *E. coli* O157:H7, C1, C2, K-12 MG1655 and lysogenic strains (C1-Φ, C2-Φ and K-12-Φ) by ELISA. The test was performed as described by Baraketi *et al*. (2018). Briefly, strains were incubated in TSB at 37°C. After O/N incubation, bacterial cultures were diluted 1/10 in TSB supplemented with mitomycin C (50 ng/mL) and were incubated at 37°C for 20 hours. The supernatants were filtered as described above. The ELISA was performed in 24-well microplates using 1 mL of each supernatant, Rabbit IgG (Sigma Aldrich, Ontario, Canada), anti-Shiga Toxin 2 as detection antibody and polyclonal anti-rabbit (Thermo Fisher Scientific, Burlington, Ontario, Canada) as secondary antibody coupled to Horseradish peroxidase (Jackson ImmunoResearch Laboratories Inc.,West Grove, PA, USA). Absorbance was read using a Biotek microplate reader at 450 nm and Gen 5 2.07 software.

## References

1. Croxen, M.A. and B.B. Finlay, Molecular mechanisms of Escherichia coli pathogenicity. Nature Reviews Microbiology, 2010. 8(1): p. 26.

2. Kaper, J.B., J.P. Nataro, and H.L. Mobley, Pathogenic escherichia coli. Nature reviews microbiology, 2004. 2(2): p. 123.

3. Wick, L.M., et al., Evolution of genomic content in the stepwise emergence of Escherichia coli O157: H7. Journal of bacteriology, 2005. 187(5): p. 1783–1791.

4. Brüssow, H., C. Canchaya, and W.-D. Hardt, Phages and the evolution of bacterial pathogens: from genomic rearrangements to lysogenic conversion. Microbiology and molecular biology reviews, 2004. 68(3): p. 560–602.

5. Canchaya, C., G. Fournous, and H. Brüssow, The impact of prophages on bacterial chromosomes. Molecular microbiology, 2004. 53(1): p. 9–18.

6. Edlin, G., L. Lin, and R. Bitner, Reproductive fitness of P1, P2, and Mu lysogens of Escherichia coli. Journal of virology, 1977. 21(2): p. 560–564.

7. Perna, N.T., et al., Genome sequence of enterohaemorrhagic Escherichia coli O157: H7. Nature, 2001. 409(6819): p. 529.

8. Hartley, M.-A., C. Ronet, and N. Fasel, Backseat drivers: the hidden influence of microbial viruses on disease. Current opinion in microbiology, 2012. 15(4): p. 538–545.

9. Landy, A., Dynamic, structural, and regulatory aspects of lambda site-specific recombination. Annual review of biochemistry, 1989. 58(1): p. 913–941.

10. Schmidt, H., Shiga-toxin-converting bacteriophages. Research in Microbiology, 2001. 152(8): p. 687–695.

11. Herold, S., H. Karch, and H. Schmidt, Shiga toxin-encoding bacteriophages – genomes in motion. International Journal of Medical Microbiology, 2004. 294(2): p. 115–121.

12. Waldor, M.K. and D.I. Friedman, Phage regulatory circuits and virulence gene expression. Current opinion in microbiology, 2005. 8(4): p. 459–465.

13. Meessen-Pinard, M., O. Sekulovic, and L.-C. Fortier, Evidence of in vivo prophage induction during Clostridium difficile infection. Applied and environmental microbiology, 2012. 78(21): p. 7662–7670.

14. Brena-Valle, M. and J. Serment-Guerrero, SOS induction by γ-radiation in Escherichia coli strains defective in repair and/or recombination mechanisms. Mutagenesis, 1998. 13(6): p. 637–641.

15. Prada Medina, C.A., et al., Survival and SOS response induction in ultraviolet B irradiated Escherichia coli cells with defective repair mechanisms. International Journal of Radiation Biology, 2016. 92(6): p. 321–328.

16. Aksenov, S.V., Induction of the SOS response in ultraviolet-irradiated Escherichia coli analyzed by dynamics of LexA, RecA and SulA proteins. Journal of biological physics, 1999. 25(2-3): p. 263–277.

17. Weisberg, R.A., et al., Role for DNA homology in site-specific recombination: The isolation and characterization of a site affinity mutant of coliphage λ. Journal of Molecular Biology, 1983. 170(2): p. 319–342.

18. Kimura, T., et al., Repression of sigK intervening (skin) element gene expression by the CI-like protein SknR and effect of SknR depletion on growth of Bacillus subtilis cells. Journal of bacteriology, 2010. 192(23): p. 6209–6216.

19. Ptashne, M. and A.G. Switch, Phage Lambda and Higher Organisms. Cell and Blackwell Scientific, Cambridge, MA, 1992.

20. Koutchma, T., L.J. Forney, and C.I. Moraru, Ultraviolet light in food technology: principles and applications. 2009: CRC press.

21. Lacroix, M. and C. Vigneault, Irradiation treatment for increasing fruit and vegetable quality. 2007: p. 1-8.

22. Gould, W.D., McCready, R. G. L., Rajan, S., & Krouse, H. R., Stable isotope composition of sulphate produced during bacterial oxidation of various metal sulphides. Biohydrometallurgy–Proceedings of the International Symposium Held at Jackson Hole, Wyoming, 1989: p. 13–18.

23. Gaougaou, G., et al., Effect of β-lactam antibiotic resistance gene expression on the radio-resistance profile of E. coli O157: H7. Heliyon, 2018. 4(12): p. e00999.

24. Nadon, C., et al., PulseNet International: Vision for the implementation of whole genome sequencing (WGS) for global food-borne disease surveillance. Eurosurveillance, 2017. 22(23): p. 30544.

25. Lu, Z. and F. Breidt, Escherichia coli O157: H7 bacteriophage Φ241 isolated from an industrial cucumber fermentation at high acidity and salinity. Frontiers in microbiology, 2015. 6: p. 67.

26. Makino, K., et al., Complete nucleotide sequence of the prophage VT2-Sakai carrying the verotoxin 2 genes of the enterohemorrhagic Escherichia coli O157: H7 derived from the Sakai outbreak. Genes & genetic systems, 1999. 74(5): p. 227–239.

27. Plunkett III, G., et al., Sequence of Shiga toxin 2 phage 933W from Escherichia coli O157: H7: Shiga toxin as a phage late-gene product. Journal of bacteriology, 1999. 181(6): p. 1767–1778.

28. Xiong, Y., et al., A novel Escherichia coli O157: H7 clone causing a major hemolytic uremic syndrome outbreak in China. PloS one, 2012. 7(4): p. e36144.

29. Welch, R., Pore-forming cytolysins of Gram-negative bacteria. Molecular microbiology, 1991. 5(3): p. 521–528.

30. Caillet, S., F. Shareck, and M. Lacroix, Effect of gamma radiation and oregano essential oil on murein and ATP concentration of Escherichia coli O157: H7. J Food Prot, 2005. 68(12): p. 2571–2579.

31. Gorden, J. and P. Small, Acid resistance in enteric bacteria. Infection and immunity, 1993. 61(1): p. 364–367.

32. Moseley, B., Ionizing radiation: action and repair. Mechanisms of action of food preservation procedures, London, Elsevier Applied Science, 1989: p. 43–70.

33. Łoś, J.M., et al., Hydrogen peroxide-mediated induction of the Shiga toxinconverting lambdoid prophage ST2-8624 in Escherichia coli O157: H7. FEMS Immunology & Medical Microbiology, 2010. 58(3): p. 322–329.

34. Huber, K.E. and M.K. Waldor, Filamentous phage integration requires the host recombinases XerC and XerD. Nature, 2002. 417(6889): p. 656.

35. Licznerska, K., et al., Oxidative stress in Shiga toxin production by enterohemorrhagic Escherichia coli. Oxidative medicine and cellular longevity, 2016. 2016.

36. Shaikh, N. and P.I. Tarr, Escherichia coli O157: H7 Shiga toxin-encoding bacteriophages: integrations, excisions, truncations, and evolutionary implications. Journal of bacteriology, 2003. 185(12): p. 3596–3605.

37. Su, L., et al., Lysogenic infection of a Shiga toxin 2-converting bacteriophage changes host gene expression, enhances host acid resistance and motility. Molecular biology, 2010. 44(1): p. 54–66.

38. Veses-Garcia, M., et al., Transcriptomic analysis of Shiga-toxigenic bacteriophage carriage reveals a profound regulatory effect on acid resistance in Escherichia coli. Applied and environmental microbiology, 2015. 81(23): p. 8118–8125.

39. Xu, X., et al., Lysogeny with Shiga toxin 2-encoding bacteriophages represses type III secretion in enterohemorrhagic Escherichia coli. PLoS pathogens, 2012. 8(5): p. e1002672.

40. Feiner, R., et al., A new perspective on lysogeny: prophages as active regulatory switches of bacteria. Nature Reviews Microbiology, 2015. 13: p. 641.

41. Yokoyama, K., et al., Complete nucleotide sequence of the prophage VT1-Sakai carrying the Shiga toxin 1 genes of the enterohemorrhagic Escherichia coli O157:H7 strain derived from the Sakai outbreak. Gene, 2000. 258(1): p. 127–139.

42. Plunkett, G., et al., Sequence of Shiga Toxin 2 Phage 933W fromEscherichia coli O157: H7: Shiga Toxin as a Phage Late-Gene Product. Journal of bacteriology, 1999. 181(6): p. 1767–1778.

43. Grandori, R., et al., Biochemical Characterization of WrbA, Founding Member of a New Family of Multimeric Flavodoxin-like Proteins. Journal of Biological Chemistry, 1998. 273(33): p. 20960–20966.

44. Uhlich, G.A., et al., Stx1 prophage excision in Escherichia coli strain PA20 confers strong curli and biofilm formation by restoring native mlrA. FEMS Microbiology Letters, 2016. 363(13): p. fnw123–fnw123.

45. Ptashne, M., & Switch, A. G, Phage lambda and higher organisms. Cell and Blackwell Scientific, Cambridge, MA., 1992.

46. Harris, D.R., et al., Directed evolution of ionizing radiation resistance in Escherichia coli. Journal of bacteriology, 2009. 191(16): p. 5240–5252.

47. Wang, X., et al., Cryptic prophages help bacteria cope with adverse environments. Nature communications, 2010. 1: p. 147.

48. Dalziel, K., Isocitrate dehydrogenase and related oxidative decarboxylases. FEBS letters, 1980. 117(S1): p. K45–K55.

49. Greener, A. and C. Hill, Identification of a novel genetic element in Escherichia coli K- 12. Journal of bacteriology, 1980. 144(1): p. 312–321.

50. Hull, A., et al., Mitomycin immunoblot colony assay for detection of Shiga-like toxin-producing Escherichia coli in fecal samples: comparison with DNA probes. Journal of clinical microbiology, 1993. 31(5): p. 1167–1172.

51. Fornelos, N., D.F. Browning, and M. Butala, The use and abuse of LexA by mobile genetic elements. Trends in microbiology, 2016. 24(5): p. 391–401.

52. Yasuda, T., et al., Physical interactions between DinI and RecA nucleoprotein filament for the regulation of SOS mutagenesis. The EMBO Journal, 2001. 20(5): p. 1192–1202.

53. de Vargas, L.M. and A. Landy, A switch in the formation of alternative DNA loops modulates lambda site-specific recombination. Proceedings of the National Academy of Sciences, 1991. 88(2): p. 588–592.

54. Fogg, P.C.M., et al., Identification of multiple integration sites for Stx-phage Φ24B in the Escherichia coli genome, description of a novel integrase and evidence for a functional anti-repressor. Microbiology, 2007. 153(12): p. 4098–4110.

55. Evans, T., R.G. Bowers, and M. Mortimer, Modelling the stability of Stx lysogens. Journal of theoretical biology, 2007. 248(2): p. 241–250.

56. Gaougaou, G., et al., Effect of β-lactam antibiotic resistance gene expression on the radio-resistance profile of E. coli O157:H7. Heliyon, 2018. 4(12): p. e00999.

57. Chen, S., et al., fastp: an ultra-fast all-in-one FASTQ preprocessor. bioRxiv, 2018: p. 274100.

58. González-Domínguez, J. and B. Schmidt, ParDRe: faster parallel duplicated reads removal tool for sequencing studies. Bioinformatics, 2016. 32(10): p. 1562–1564.

59. Bankevich, A., et al., SPAdes: a new genome assembly algorithm and its applications to single-cell sequencing. Journal of computational biology, 2012. 19(5): p. 455–477.

60. Latif, H., et al., A gapless, unambiguous genome sequence of the enterohemorrhagic Escherichia coli O157: H7 strain EDL933. Genome announcements, 2014. 2(4): p. e00821–14.

61. Darling, A.E., B. Mau, and N.T. Perna, progressiveMauve: multiple genome alignment with gene gain, loss and rearrangement. PloS one, 2010. 5(6): p. e11147.

62. Seemann, T., Prokka: rapid prokaryotic genome annotation. Bioinformatics, 2014. 30(14): p. 2068–2069.

63. Li, H. and R. Durbin, Fast and accurate short read alignment with Burrows–Wheeler transform. bioinformatics, 2009. 25(14): p. 1754–1760.

64. Li, H., et al., The sequence alignment/map format and SAMtools. Bioinformatics, 2009. 25(16): p. 2078–2079.

65. Yu, L.H., et al., Arabidopsis EDT 1/HDG 11 improves drought and salt tolerance in cotton and poplar and increases cotton yield in the field. Plant biotechnology journal, 2016. 14(1): p. 72–84.

